# SPECHTacle: An Efficient Detection and Visualization of Cell Specific Regulation in Single Cell-Based Experiments Using a Novel Combinatorial Computational Tool

**DOI:** 10.64898/2026.11.04.764311

**Authors:** Kathleen M. Specht, Hana C. Schwierling, Matthew J. Robson

## Abstract

Herein, we describe SPECHTacle, a novel computational tool affording the ability to efficiently visualize and delineate the specific cellular population(s) impacted by an experimental paradigm utilizing heterogeneous cell-based samples. SPECHTacle can be employed to efficiently determine, in a cell specific manner, the populations preferentially impacted by treatments, drugs, injuries, disease states, and/or development within datasets continuously being generated across nearly countless subfields in molecular biology. SPECHTacle accomplishes this task by harnessing the power of Waddington optimal transport analysis and Shannon entropy in a combinatorial fashion to distinguish cell populations preferentially altered due to nearly any experimental paradigm and generates a visual representation of cellular entropy (i.e. change) across entire datasets. Ultimately, this allows for efficient determination of specific populations preferentially impacted by an experimental paradigm and identifies populations to target for downstream analyses employing quantitative measurements. We propose that SPECHTacle represents an immediately useful computational tool with the potential to be integrated into single cell-based workflows and greatly enhance the efficiency of both manual and machine-based learning workflows.

## INTRODUCTION

The advent of single cell genomics technology has revolutionized our ability to characterize and define the molecular status of individual cells within complex, cellular heterogenous organ systems, such as the brain, with a high level of fidelity. Single cell (scRNA-seq) and single nucleus RNA sequencing (snRNA-seq), combined with spatial transcriptomics is becoming a versatile and widespread tool^1^ to study cell-specific landscapes^2^ in development,^3^ physiologic processes, and disease reprogramming.^4–6^ Although these types of studies are inherently powerful in their ability to discern cell specific transcriptional regulation, it has historically been difficult to determine precisely where to direct initial cell specific analyses occurring between defined experimental endpoints.^7^ To date, the bulk of these forms of studies are conducted with a preconceived notion of which specific cell populations to direct analysis procedures with regards to cell specific effects for a given experimental paradigm.^8^ Alternatively, much work has been centered upon the identification of novel cell populations in heterogeneous samples and the generation of physiologic atlases.^9–14^ While this has admittedly greatly increased our knowledge base of cell specific regulation in highly heterogenous environments, it may not identify preferentially impacted cell populations within a given experimental paradigm and/or disease state.

With these advancements, computational and machine-learning approaches have been developed to enhance information gained from these datasets.^15–17^ Waddington optimal transport (Waddington-OT or WOT) is a conceptual framework utilized to infer how individual cells/nuclei evolve with a high degree of temporal resolution.^3^ WOT is an interpretive analysis that applies the mathematical concept of optimal transport to infer the temporal coupling of transcriptional regulation occurring within defined populations between timepoints.^3^ Conrad H. Waddington (1905-1975) was a British biologist who changed the way we think about developmental biology,^18^ and is known for his contributions to the epistemology of cellular fate differentiation.^3^ The conceptual framework of WOT comes from combining CH Waddington’s contributions with the mathematical concept of optimal transport (OT). OT was originally developed to calculate the most efficient manner in which to redistribute earth and was first applied by Napoleon in Egypt.^19^ It was later generalized to identify optimal coupling of probability distributions using linear programming.^20^ WOT modifies this conservation of mass to instead account for transcriptomic regulation occurring between defined experimental timepoints.^3^

Perhaps the most useful application of WOT is to infer long-range temporal couplings from calculated couplings between consecutive timepoints.^21,22^ In other words, WOT computes individual transport matrices which infer overall changes in a cell’s fate over time. WOT can be used to assess trends towards a specific cell fate by computing a cell’s likely ancestors from an earlier timepoint, and its likely descendants in a later timepoint.^21^ This can be visualized with fate matrices and transition tables.^21,22^ This application becomes inherently limited in investigations of unknown trajectories or disease/injury states, where visualizing individual cell fate transitions becomes overly complicated and inefficient.^23^

Within the current manuscript, we build upon known, defined WOT analyses utilized across the snRNA-seq and scRNA-seq field,^24^ by calculating the Shannon entropy and creating a composite image summarizing how much or how little a cell’s fate (i.e. amount of “cellular change”) changes over time. Shannon entropy describes the amount of information that can be contained within a variable.^25,26^ Herein, we utilize individual cells/nuclei as the variable that carries information as a collection of possible fates. A cell with low entropy carries few possible fates, meaning its fate (state) is consistent over time; a cell with high entropy carries many possible fates (states), meaning that its fate is variable.^26–28^ We have dubbed this tool “SPECHTacle,” an acronym for **S**tochastic **P**rojection of **E**ntropic **C**ell **H**istory and **T**ransition; that is representative of the profound visuals generated through the analysis of divergent populations of cells/nuclei. SPECHTacle is a visualization tool that utilizes fate entropy calculations to construct composite color-scaled Uniform Manifold Approximation and Projections (UMAPs) reflecting the levels of entropy in each individual datapoint, specific cell and/or nuclei.^29^ SPECHTacle allows for the visualization of overall variations in cell fates/transcriptional alterations as compared to initial and/or control populations. SPECHTacle thereby affords the rapid visualization of overt transcriptional regulation over time across diverse cell populations and/or defined cell specific subpopulations captured, representing a novel and useful tool to those engaged in delineating developmental and/or cellular regulation over time, due to disease states, drug treatments or nearly any other experimental paradigm in either preclinical or clinically obtained datasets.

## METHODS

### Runx1*^+^ Dataset*

The SPECHTacle paradigm was exemplified using the publicly available data set GSE224407 accessed through the Gene Expression Omnibus (GEO) from the National Center for Biotechnology Information. It was originally published under the title “Single-Nucleus RNA Sequencing of Developing and Mature Superior Colliculus Identifies Neuronal Diversity and Candidate Mediators of Circuit Assembly” by Choi et al. in 2023.^30^ This study involved the isolation and sequencing of nuclei from the superior colliculus of C57BL/6J mice on embryonic day 19 (E19) and post-natal days 4, 8, and 21 (P4, P8, and P21, respectively). The nuclei were sequenced using the 10x Genomics 3’ v3 snRNA-seq platform.^30^

Original raw .h5 files were downloaded from the GEO database and then uploaded into Spyder (v. 6.1.0) with Python (v. 3.10). Spyder was run in a custom-made conda environment (dependencies of the environment can be found in Supplemental Table 1). The data was prepared into one AnnData object. To first test SPECHTacle on a specific cell population, the object was subset to only include cells that exhibited positive expression of *Runx1*, a gene used in the original analysis by Choi et al. to mark microglial cells. Using scanpy (Single-Cell Analysis in Python, v. 1.11.5)^31^ the data was normalized and scaled before running principal component analysis (PCA) and the find neighbors function. Finally, a UMAP (a 2D depiction of cells in PCA space) was produced. All plots were produced using matplotlib.pyplot (matplotlib v 3.10.8). The object was then saved as a .h5ad file.

This .h5ad file was then loaded into the SPECHTacle script as the adata_file. The workflow was tailored to this dataset using the parameters listed in Supplemental Table 2. These can be tailored to the user by altering these parameters denoted by ‘’’ P‘’’ in the code. The PCA, neighbors graph, and UMAP were rerun to ensure they were stored after the loading. Next, Leiden clusters were computed with a relatively low resolution of 0.3 to optimize the subclusters generated while preserving visual differences between them.

The model assumed E19 (the earliest timepoint available within the dataset) as the starting point and designated it as numerical time 0. Subsequently, P4 nuclei were designated as numerical time 4; P8 as numerical time 8, and P21 as numerical time 21. Here, numerical time serves as a discrete variable denoting the real time at which samples came from. Numerical time was then translated to a categorical timepoint variable, where numerical time 0 serves as the starting timepoint (T0), while numerical time 21 is the terminal timepoint (T3). Intermediate numerical times 4 and 8 respectively become T1 and T2. Thus, numerical times 0, 4, 8, and 21 become the respective timepoint variables T0, T1, T2, and T3.

To assign samples to these timepoint variables, we first used indicators embedded in sample file names to sort samples into a categorical sample variable, where any file indicated as E19 was assigned to the sample variable S1. Similarly, P4 files were assigned to S2, P8 to S3, and P21 to S4. These sample variables were then matched to their respective timepoint variables, so E19 files assigned to sample variable S1 are associated to the corresponding timepoint variable T0. P4/S2 is associated to T1, P8/S3 is associated to T2, and P21/S4 is associated to T3. These sample/timepoint associations are stored as appropriate metadata.

The OT model (from wot.ot import OTModel, wot v 1.0.8.post2) utilized the following parameters for transport map computations: = 0.1, λ_1_ = 3.0, and λ_2_ = 80.0. represents the entropic regularization parameter that determines the “spread” of cell fates, where larger values give more descendant distributions ( = 0.1 is moderate).^3,21^ λ represents the growth regularization parameters of the starting (λ_1_) and terminal (λ_2_) points, thus controlling the strictness of ancestor-descendant couplings with larger values enforcing strict constraints (λ_1_ = 3.0 is moderate; λ_2_ = 80.0 is high).^3,21^ Thus, our model utilized a moderate degree of entropy and strictly enforced temporal couplings.

Transport maps were individually created for each transition, including T0 → T1, T0 → T2, T0 → T3, T1 → T2, T1 → T3, and T2 → T3. Transport maps are computed by calculating the amount of “mass” moving from timepoint to timepoint based on the cost-efficiency of moving that PCA distance.^3,21^ The individual transport maps were transformed into matrices (pandas v 2.3.3). A stepwise transport map was then calculated by taking the dot product of the major transition matrices, including T0 → T1, T1 → T2, and T2 → T3. Cell-cell transport was computed for all matrices while preserving the original timepoint metadata and the inter-timepoint cluster couplings. Cluster fate probabilities were calculated before proceeding with Shannon entropy calculations (from scipy.stats import entropy, scipy v 1.15.3).^32^ Computed entropy values were stored as fate entropy scores that are then projected onto a corresponding, composite UMAP. The T0 cells were plotted on the composite UMAP scatterplot and colored based on their fate entropy scores from the stepwise calculations. The Leiden clusters were then overlayed on that plot as cluster outlines (numpy v 2.2.6).

### Psilo-Seq Dataset

Next, we wanted to evaluate and validate the utility of SPECHTacle in a large preclinical dataset that employed a treatment vs control setting. To do this, we used the publicly available dataset BioProject ID PRJNA1204073 (nicknamed Psilo-seq), from the work titled “Single-nucleus transcriptomics reveals time-dependent and cell-type-specific effects of psilocybin on gene expression” by Liao, O’Farrell, Weiner, et al. posted in 2026^33^ (in pre-print at the time of this analysis). Liao et al. randomly assigned C57BL/6J mice to psilocybin or ketamine treatment groups and collected samples from the dorsal medial frontal cortex at 1, 2, 4, 24, or 72 hours post-injection. Non-drugged, age-matched mice were used as controls. For the sake of simplicity in our analysis, we used only the psilocybin or control treated samples from 1, 24, and 72 hours in our analysis. Publicly available data was downloaded and prepared as described above.

To assess psilocybin-induced changes, we used control samples as the starting point or numerical time 0, with 1-hour, 24-hours, and 72-hours post-psilocybin samples becoming numerical times 1, 24, and 72 (respectively). Following the same categorial timepoint association method as before, numerical time 0 becomes T0, numerical time 1 becomes T1, numerical time 24 becomes T2, and numerical time 72 becomes the terminal T3. Sample-timepoint associations were created as before, with control samples becoming S1 to be associated with T0. Following this framework, 1-hour post-psilocybin samples became S2, 24-hour samples become S3, and 72-hour samples become S4. Finally, control/S1 samples were associated with T0, 1-hour/S2 with T1, 24-hour/S3 with T2, and 72-hour/S4 with T3. WOT and SPECHTacle analyses were conducted as before. An additional cell-type workflow was utilized here to test SPECHTacle on data with diverse cell populations, where Shannon fate entropy calculations were visualized by both Leiden cluster and cell type metadata publicly available and associated with the dataset.

## RESULTS

### SPECHTacle Output

Running the SPECHTacle specific-code outputs 10-12 .png image files, depending on whether the additional cell type entropy visualization is included. The figures appear in the code in the same order they are listed and described here. umap_leiden_wot.png shows the Leiden clustering used for the WOT model with included cluster legend as in **Figures 1A and 4A**. umap_timepoint.png shows the same Leiden clustered UMAP with cells colored by timepoint rather than cluster, as in **Figures 1B and 4B**. Timepoint legend is included in the image. The next six images are SPECHTacle projections showing Shannon fate entropy scores in individual comparisons, as in **Figures 2A-F and 5A-F**: fate_entropy_t0_t1_umap.png, fate_entropy_t0_t2_umap.png, fate_entropy_t0_t3_umap.png, fate_entropy_t1_t2_umap.png, fate_entropy_t1_t3_umap.png, fate_entropy_t2_t3_umap.png. Next, fate_entropy_transitionary_umap.png shows a final SPECHTacle projection showing overall fate entropy calculated from the stepwise transport map (as in **Figures 2H and 6A**). This UMAP is displayed as a scatterplot in entropy_with_cluster_oultines_leiden.png (as in **Figures 3 and 6B**). Here, each cluster is outlined according to its Leiden cluster identity (a cluster legend is not included in the raw image). The remaining two images are only included in the cell type entropy visualization. umap_celltype.png shows the same cells labeled with cell type metadata, rather than the Leiden cluster (with included cell type legend), as in **Figure 7A**. umap_fate_entropy_celltype_outlines.png shows a scatterplot of fate entropy with cell type colored-outlines (a cell type legend is included in the raw image), as in **Figure 7B**.

**Figure 1:**
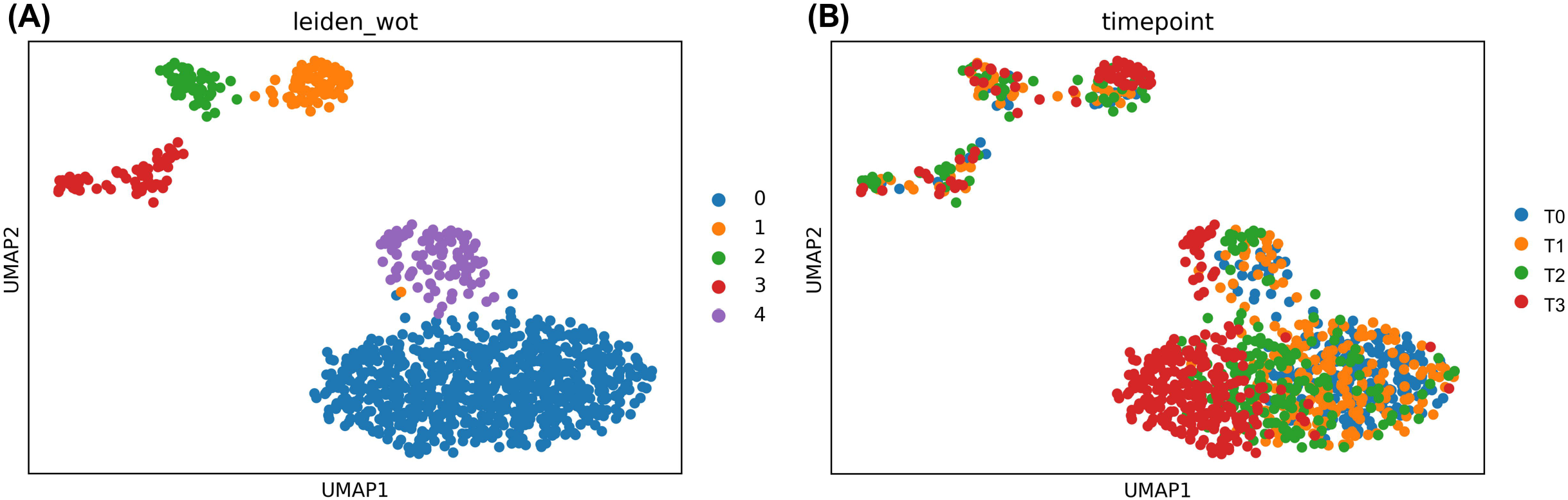
Leiden Clustering of *Runx1*^+^ Nuclei. (A) UMAP depicting identifiable, distinct clusters of the *Runx1*^+^ population within the dataset. (B) UMAP representation of *Runx1*^+^ nuclei by timepoint. Starting timepoint contains nuclei collected on embryonic day 19 (E19), referred to as T0. Remaining post-natal timepoints contain nuclei collected on post-natal days 4, 8, and 21, respectively, whereby P4 nuclei makeup T1, P8 nuclei makeup T2, and P21 nuclei makeup T3. The timepoints represented within *Runx1*^+^ nuclei mix relatively well, with some separation noted of the T3 nuclei (P21).

**Figure 2:**
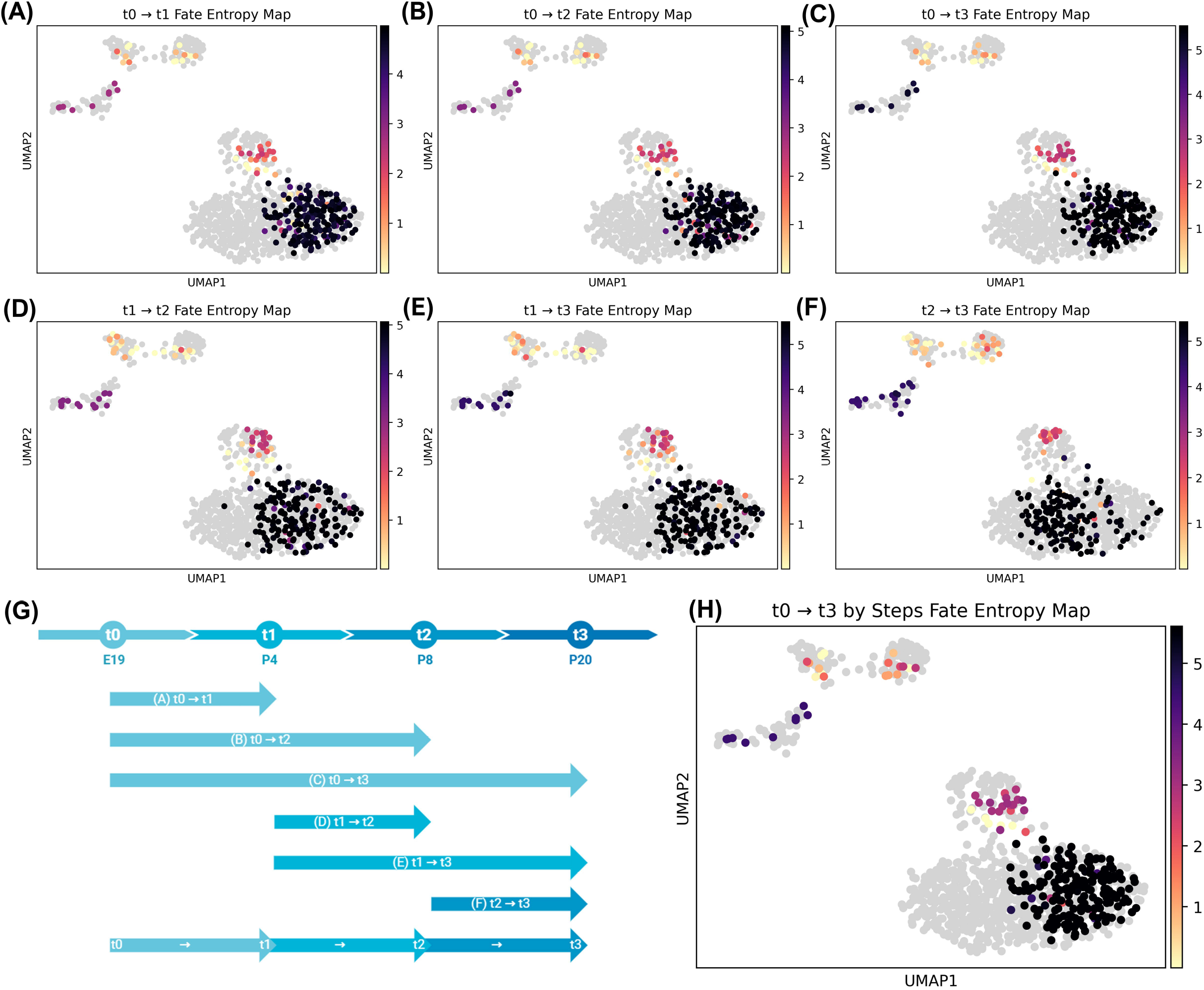
SPECHTacle Projections Showing Entropy Calculated at Each Temporal Transition. (A-F) UMAPs representing levels of Shannon entropy calculated at individual transitions, corresponding to timeline depiction (G). Colored cells represent the starting fate at each point, while grey cells represent the ending fate at each point. WOT + SPECHTacle visualization predicts how the starting cells transition into the ending cells, so only starting cells are colored to show their predicted levels of entropy. (A-C) Colored T0 cells represent predicted change occurring within these nuclei between T0 → T1 (A), T0 → T2 (B), and T0 → T3 (C). (D, E) Colored T1 cells represent predicted change occurring between T1 → T2 (D) and T1 → T3 (E). (F) Colored T2 cells represent change predicted to occur between T2 → T3. Darker coloration represents higher levels of calculated cellular entropy, meaning they undergo high levels of change during the corresponding transition; lighter colored cells have less calculated entropy, i.e. less change. (H) Final UMAP colored by terminal fate entropy computed from individual transport maps. This projection infers the total overall entropy calculated from cells across all timepoints, T0 → T1 → T2 → T3.

**Figure 3:**
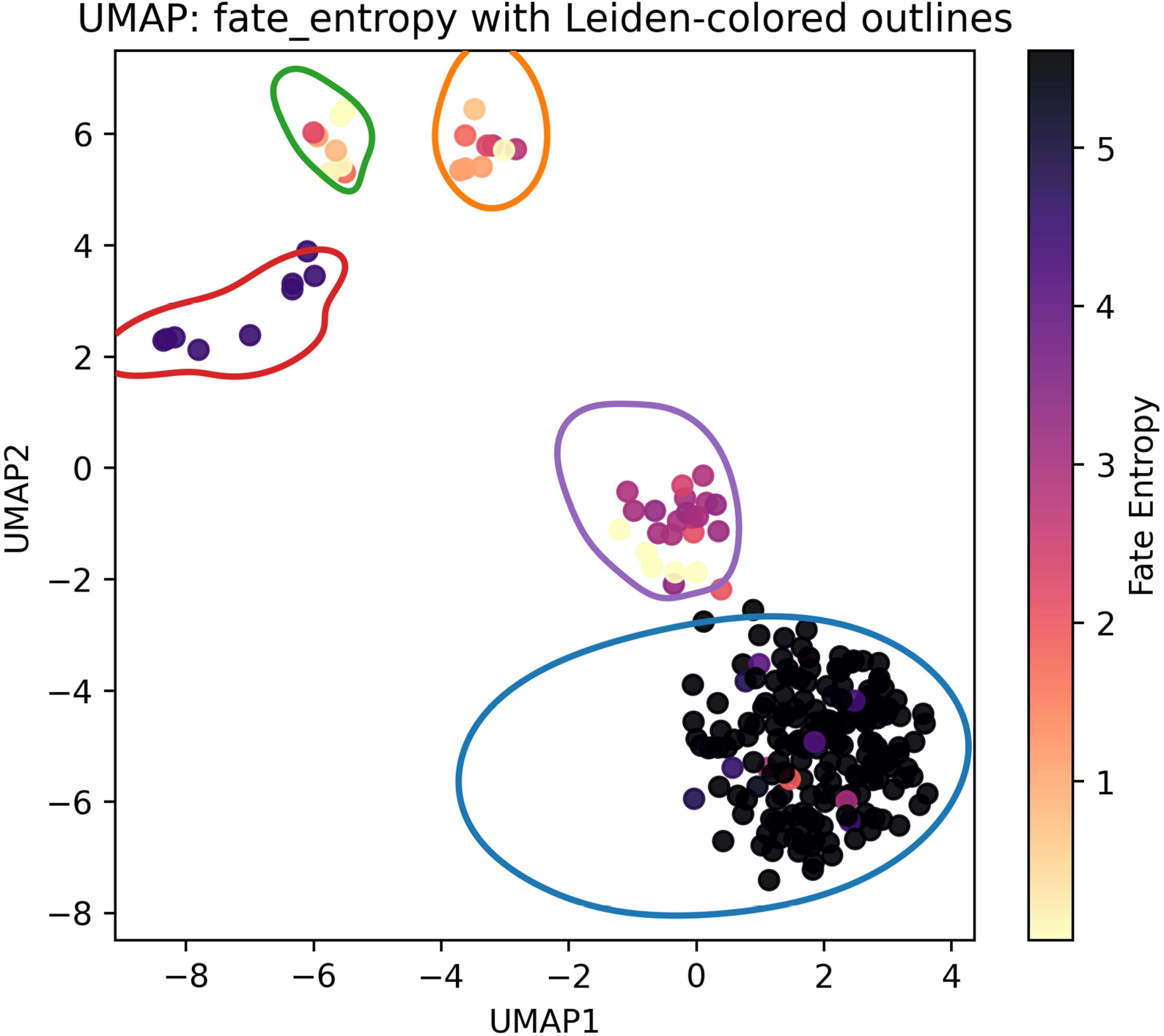
SPECHTacle Projection with Cluster Outlines. UMAP with T0 cells colored by Shannon Entropy scores based on a transport map-built transport matrices between T0 → T1, T1 → T2, and T2 → T3. Colored polygons identify and mark original Leiden cluster identities. Cluster 0 clearly has the highest calculated cellular entropy, while Clusters 1 and 2 have the lowest. Clusters 3 and 4 were calculated to have intermediate levels of entropy. Entropy rankings across the identifiable populations of *Runx1*^+^ nuclei demonstrate how SPECHTacle can be used to identify sub-clusters within a particular cell type exhibiting a high level of cellular entropy across timepoints represented within a dataset.

**Figure 4:**
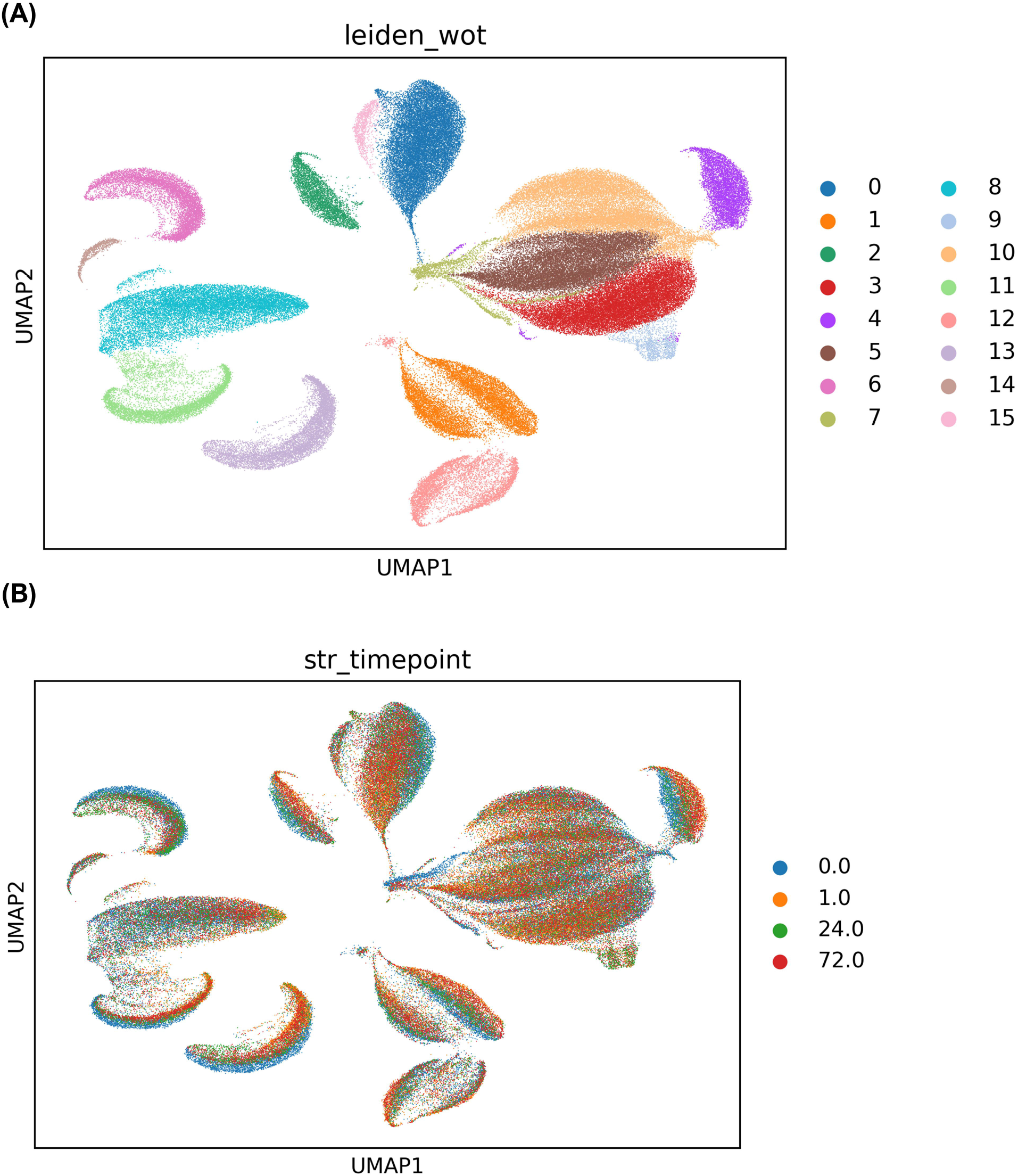
Leiden Clustering of Diverse Cell Populations from Brains of Psilocybin Dosed Mice. (A) UMAP depicting identifiable nuclei clusters from samples isolated from mice treated with either vehicle or psilocybin. Clustering was conducted using a resolution of 0.3 to generate depicted UMAP. (B) UMAP showing distribution of nuclei by timepoint. Starting timepoint contains nuclei from control mice (blue), referred to as T0, and is considered 0 hours post-injection (hpi). Remaining timepoints contain nuclei collected 1, 24, or 72 hpi of psilocybin. The nuclei collected after 1 hour makeup T1 (orange), 24 hpi makeup T2 (green), and 72 hpi makeup T3 (red). Combined, identifiable nuclei populations are homogenously distributed across timepoints, with minimal stratification of Cluster 6.

### *Developmental Analysis of* Runx1*^+^ Nuclei*

We first assessed SPECHTacle on a subset of the *Runx1*^+^ microglia. The Leiden clustering plot **(Figure 1A)** specifically shows determined groupings of cells with similar gene expression, representing specific, definable subpopulations of *Runx1^+^* nuclei. At a resolution of 0.3, this *Runx1*^+^ population is split into 5 definable subpopulations. The same base UMAP is utilized to examine distribution across timepoints **(Figure 1B)**. It should be noted that appropriate quality controls, including assertions of even cellular/nuclei distributions across timepoints is a requirement for the inherent validity of analysis. If timepoints cluster separately, it can become difficult to accurately interpret temporal couplings between timepoints. Examination of the UMAP shows fair distribution in the sample population, with Cluster 0 exhibiting some stratification of P21 nuclei, that inherently represents the transitional state of the cell population at hand.

Once fate entropy scores were calculated for all transitions, the scores could be plotted on the UMAP. Looking at the first transition from T0 → T1 **(Figure 2A)**, it is evident that the entropy is not uniform across the cells. Here, T0 cells are colored while all other cells are grey, signifying the predicted level of entropy displayed by T0 cells during their transition to T1. T0 cells in the bottom cluster (Cluster 0) exhibit the highest entropy, therefore being the most likely to undergo a change in fate identity. A similar trend holds in the transition from T0 → T2 **(Figure 2B)**, where it is clear Cluster 0 again has the most entropy. From T0 → T3 **(Figure 2C)**, this trend is maintained with slightly higher entropy in other clusters, including the cluster on the left (Cluster 3).

The intermediate transition states can be examined as well. In visualizing T1 → T2 **(Figure 2D)**, T1 cells are now colored with similar cluster entropy trends. The T1 → T3 **(Figure 2E)**, indicates that this trend is maintained over time. The latest transition from T2 → T3 **(Figure 2F)** (with T2 cells colored) further shows a clear entropy trend later in the timeline. A visual representation of this timeline and the transitions **(Figure 2G)** aids in interpreting which plots correspond to which timepoints.

To paint the clearest picture of the changes occurring from start to finish, the timeline can be broken down in stages. Considering only the major transitions T0 → T1, T1 → T2, and T2 → T3 gives a comprehensive picture of what is happening between the beginning and terminal fates. To plot this, each transport matrix is taken together to compute the total entropy for T0 → T1 → T2 → T3 **(Figure 2H)**. This total entropy calculation can be used to sum the entropy trends of each stage. As expected, Cluster 0 exhibits the highest entropy, followed by Clusters 3 and 4 with intermediate entropy, and Clusters 1 and 2 with the lowest entropy. This composite/stepwise entropy analysis is plotted with original Leiden cluster coordinates as outlines **(Figure 3)**, allowing for the assessment of inter-cluster differences in entropy levels. Thus, highly dynamic subpopulations can be delineated, and their levels of transcriptional regulation in response to experimental paradigms can be visualized over time.

### Treatment Effect Analysis on Psilocybin vs Control Treated Cells

Running SPECHTacle on a subset of the Psilo-seq dataset enabled assessment of its utility in larger single cell and single nucleus datasets with many cell types as well as to evaluate transcriptomic changes induced by biologic perturbation (i.e. psilocybin treatment). Inputting the dataset into our pipeline revealed 16 clusters **(Figure 4A)**. Non-drugged control samples were designated as T0 to serve as our baseline population for the investigation of psilocybin-induced changes. Psilocybin-treated samples from the other time points, including T1 (1 hour post-injection), T2 (24 hour post-injection), and T3 (72 hour post-injection) were dispersed evenly across clusters, an effect that enhances the ability of SPECHTacle to confidently identify applicable cell populations and their respective entropy as noted above **(Figure 4B)**. The analysis revealed heterogenous levels of entropy across clusters **(Figure 5)**, with clusters 0, 3, 5, and 10 having the highest entropy scores. Thus, these clusters underwent the most change from baseline after psilocybin treatment, indicating that the transcriptomes of these cells were largely impacted by treatment **(Figure 6)**.

**Figure 5:**
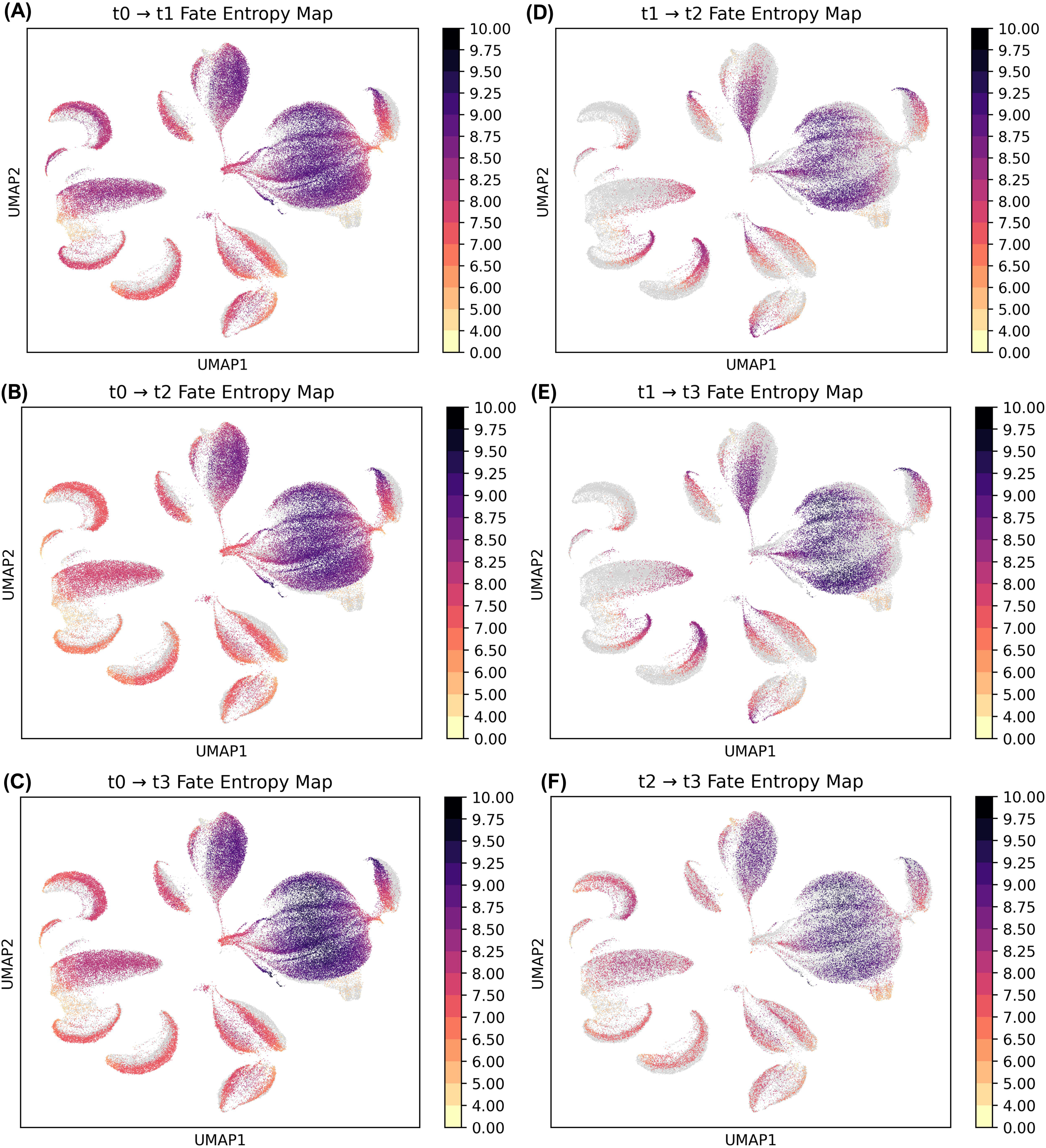
SPECHTacle Projections Visualizing Calculated Cellular Entropy Across Timepoint. (A-F) UMAPs representing calculated levels of Shannon entropy at individual transitions, similar to that depicted within Figure 2. Darker coloration represents higher levels of calculated cellular entropy, indicative of greater levels of change; lighter coloration represents less calculated cellular entropy, meaning these specific populations exhibit less change in response to treatment. (A-C) Colored T0 cells represent predicted change occurring between T0 → T1 (A), T0 → T2 (B), and T0 → T3 (C). (D, E) Colored T1 cells represent predicted change occurring between T1 → T2 (D) and T1 → T3 (E). (F) Colored T2 cells represent change predicted to occur between T2 → T3.

**Figure 6:**
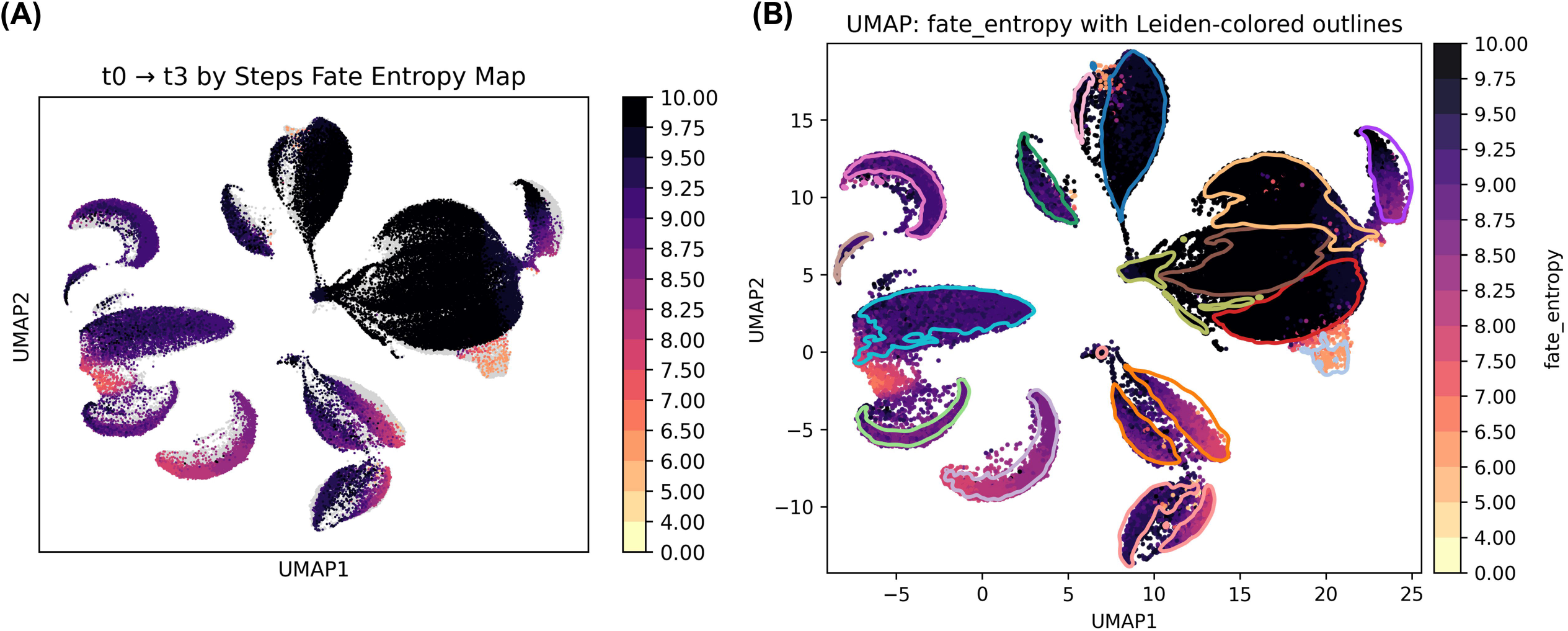
SPECHTacle Projection with Leiden Cluster Outlines of Psilocybin Injected and Control Mice. (A) Final UMAP with coloration depicting terminal fate entropy computed from individual transport maps depicted within Figure 5. UMAP projection conveys overall cellular entropy of identifiable populations across all timepoints, from T0 → T1 → T2 → T3. (B) Provided outlines identify original Leiden cluster identities within the dataset. Clusters 0, 3, 5, and 10 show the highest levels of entropy, with Clusters 9 and 13 exhibiting the lowest calculated levels of cellular entropy as a result of psilocybin administration. Visualization using SPECHTacle in this manner, allows for the rapid identification of cellular populations most impacted by an experimental paradigm, in this instance, the administration of a single dose of psilocybin to murine subjects.

The original analysis by Liao et al. included cell annotations stored as metadata. These annotations were used to color the original Leiden-clustered UMAP **(Figure 7A)**. SPECHTacle was designed to conduct an additional cell type analysis if such metadata exists and is specified. To visualize this, cell type outlines were plotted over the stepwise entropy scatterplot **(Figure 7B)**. This allows for the stratification of cell types by entropy level. Here, the highest entropy was observed among identifiable populations of glutamatergic neurons, particularly among intratelencephalic glutamatergic identities as defined in the original analysis 004 L6 IT CTX Glut, 005 L5 IT CTX Glut, 006 L4/5 IT CTX Glut, and 007 L2/3 IT CTX Glut. All other cell types displayed comparatively lesser entropy, including GABA neurons (046 Vip Gaba, 047 Sncg Gaba, 049 Lamp5 Gaba, 052 Pvalb Gaba, and 053 Sst Gaba), astrocytes (319 Astro-TE NN), oligodendrocytes/precursors (327 Oligo NN and 326 OPC NN), endothelial cells (333 Endo NN), and microglia (334 Microglia NN).

**Figure 7:**
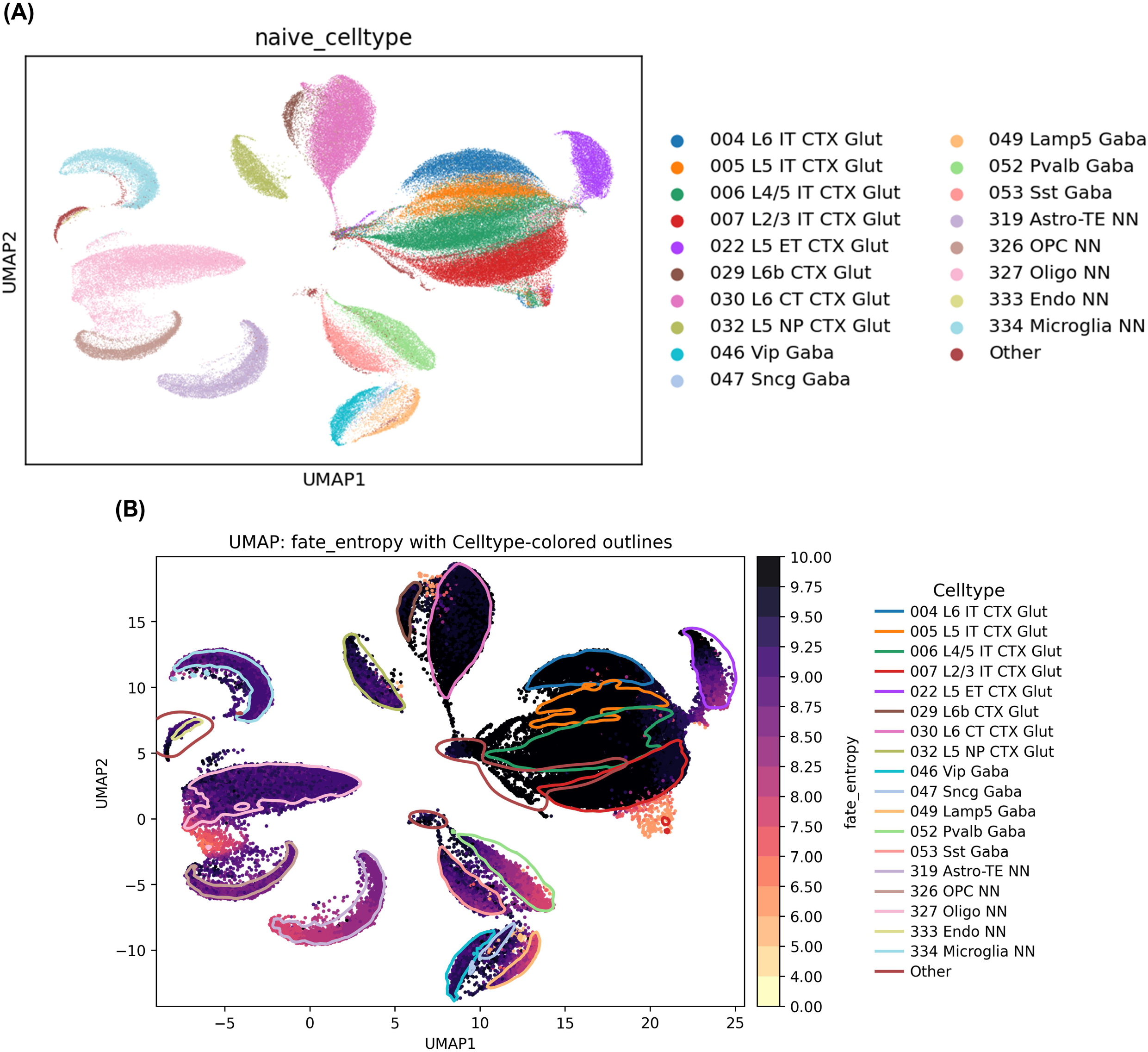
SPECHTacle Projection with Celltype Cluster Outlines of Psilocybin Injected and Control Mice. (A) UMAP depiction of divergent cell populations captured within samples, as determined by Liao et. al. (B) Provided outlines mark identifiable populations of nuclei from full dataset. SPECHTacle analysis revealed that glutamatergic neuron populations exhibit the highest calculated levels of cellular entropy, whereas oligodendrocytes and oligodendrocyte precursor cells (OPCs) have the lowest levels of calculated cellular entropy. Visualization of the dataset using SPECHTacle allows for the rapid assessment that glutamatergic neuron populations exhibit a high level of overt change induced by psilocybin administration, as compared to other identifiable populations.

## DISCUSSION

Herein, we present and provide to the greater research community, a novel analysis methodology, dubbed SPECHTacle. SPECHTacle is a combinatorial, computational tool useful in both analyzing and subsequently visualizing transcriptomic change and alterations within single cell and single nucleus RNA sequencing datasets using a novel combination of Waddington Optimal Transport (WOT) and Shannon entropy calculations. SPECHTacle enables high fidelity visualization of UMAP-structured data to model cell fates determined by transcriptional regulation over time through UMAP-projections of WOT + Shannon Fate Entropy, thus allowing for overt visualization of predicted change occurring in defined cell population(s).

Overall, our novel workflow has potential utility in any scRNA-seq/snRNA-seq investigation with or without a time variable. SPECHTacle can be applied to endless areas; from drug response, injury and disease states to developmental and evolutionary research. SPECHTacle was designed as a versatile tool allowing users to tailor their analysis paradigm based on the context of their investigation; whether that is establishing developmental timelines or identifying highly variable cell types in a disease model. Lastly, SPECHTacle is capable of quickly and efficiently identifying cellular targets impacted by drug treatment, injury, disease, or cellular development over time to allow for informed experimental validation.

SPECHTacle enables visual and quantitative distinction between cells of rigid fate and cells of more variable fate induced by nearly any variable or experimental paradigm. Rigid fate is marked by lower entropy and indicates that a cell is unlikely to have a terminal fate different from its beginning fate, moving very little in PCA space. Variable fate is marked by higher entropy and the potential to move farther in PCA space from beginning to terminal fate. Fate determinations are important in cell development allowing for the stratification of progenitor and mature cell types and/or cellular regulation occurring within defined disease states.

To test the utility of SPECHTacle, we employed the computational analysis on two distinct publicly available datasets. First, the GEO dataset GSE224407 generated by Choi et al. was used to exemplify SPECHTacle in the context of a small, defined cell population in a developmental setting.^30^ Namely, a subset of *Runx1*^+^ nuclei from the superior colliculus was run through the SPECHTacle pipeline to reveal varying levels of entropy within the subclusters. Due to the focus on developmental timepoints, SPECHTacle uncovered those populations were still undergoing maturation processes during postnatal periods contained within the dataset. Cluster 3 nuclei were a population identified as having increased cellular entropy at each transition. The highest entropy was observed between T2 → T3, corresponding to post-natal days 8 and 21, respectively. This suggests that post-natal days 8 through 21 are a critical period of development within this *Runx1^+^* subpopulation. While the identification of the particular subpopulations is beyond the scope of the current manuscript, it is subsequently possible to identify the unique differences between subpopulations with the highest entropy. For example, cluster 1 may represent a homeostatic microglia population, exhibiting a low level of overt change and therefore a low level of entropy. The precise changes driving differences in cellular entropy are also beyond the scope of the current manuscript, as our analyses solely exemplify the utility of the workflow in investigating the development of the particular cell types and subpopulations.

Second, the Psilo-Seq dataset (BioProject ID PRJNA1204073) generated by Liao et al.^33^ exemplified SPECHTacle’s utility in identifying targetable cell types within larger datasets. This analysis also served to demonstrate how SPECHTacle may be used to classify cell types most affected by an experimental perturbation such as drug treatment, injury, or disease-state. Here, SPECHTacle uncovered intratelencephalic glutamatergic neurons as the most highly impacted cell population over the first 72 hours after psilocybin-treatment. Although identifying the specific molecular mechanisms associated with cell-specific alterations is beyond the scope of the current tool, SPECHTacle can be useful in the optimization of workflows to identify specific cell populations and their definable subpopulations to target for the identification of applicable molecular pathways driving biological phenotypes. It is in this manner that SPECHTacle represents an extremely powerful hypothesis generating tool from a myriad of previously generated and future datasets.

Additionally, the original analysis comparing control and psilocybin treated animals performed by Liao et. al. concluded that the greatest number of differentially expressed genes (DEGs) was found at 1 hour after injection, almost no DEGs at 24 hours, and a resurgence of DEGs again at 72 hours.^33^ Our analysis indicates a moderate amount of entropy during the first transition (T0 → T1; control → 1hr) **(Figure 5A)**. We found the lowest entropy scores for the second transition (T0 → T2; control → 24 hours) consistent with the lack of DEGs found in the original analysis at this timepoint (**Figure 5B**). While both analyses are consistent with each other an entropy analysis captures different information than solely accounting for the sheer number of DEGs. While DEGs indicate changes in the overall gene expression value for individual transcripts within definable cell populations, cellular entropy indicates changes in the overall functions of the cell taking into account gene distribution. In this way, cellular entropy takes into account the information content and heterogeneity of the transcriptome and not solely the number of DEGs.^27,34^ This means it is possible to have transitions that have low entropy scores and high amounts of DEGs and high entropy scores with low DEGs. While the high number of DEGs between 0 and 1 hour informs us that there are a lot of changes in gene expression levels, the high entropy between 0 and 72 hours informs the user that there is a high level of change in cellular function occurring during that time.

Perhaps the most useful application of this tool comes from the ability of SPECHTacle to use control samples as the baseline population (designated as numerical time 0 or T0), allowing for comparisons between treatments and across time. Drug-treated or otherwise perturbed samples collected at progressing time intervals can serve as remaining timepoints and terminal fate, to create a pseudo-time fate trajectory analysis. Alternatively, SPECHTacle could be applied in the context of before and after treatment without timepoint information to identify cell targets that are most affected by experimental intervention. If cell annotations are included in the loaded dataset, SPECHTacle can also conveniently compare diverse cell types within the data to identify which cell types exhibit the greatest amount of functional alterations; therefore, streamlining the discovery of preferentially impacted populations that may have been previously hypothesized to be unaffected.

The primary limiting aspect of employing SPECHTacle is the requirement of even distribution of cells across timepoints/experimental variables. The WOT conceptual basis of SPECHacle predicts temporal couplings of cells, leaving room for bias if timepoints cluster separately. We observed this kind of separation in our developmental analysis of *Runx1*^+^ nuclei of the Choi et al. dataset, where Cluster 0 exhibits timepoint gradation. Such timepoint stratification creates ambiguity as it can indicate potential batch effects or developmental maturation. If batch effects contribute to timepoint stratification, entropy scores may become overinflated because of artificial differences imposed. Whereas developmental datasets inherently contain shifts in cell fate, which represent expected high levels of entropy. In this case, SPECHTacle can still provide context and understanding to stratified datasets; such as in the *Runx1*^+^ dataset, where it appears changes due to developmental maturation contribute to the timepoint stratification. However, it is beyond the scope of this paper to make a conclusive determination whether batch effects are an additional contributing factor.

In contrast, our analysis of the Psilo-Seq dataset by Liao et al. had a more even timepoint distribution **(Figure 4B)**. The even distribution within these clusters prevents ambiguity caused by timepoint stratification imposed by external sources (i.e. batch effects). This underscores the importance of visualizing the UMAP by timepoint in the beginning stages of SPECHTacle to ensure validity and a thorough understanding of analysis.

To conclude, SPECHTacle provides a useful visual computational tool for molecular biologists seeking to visualize cell-specific effects in highly heterogeneous environments. Thus, the versatility of this tool enables the efficient identification of highly variable cell populations that may serve as targets for further experimental investigations and/or therapeutic discovery and development.

## DATA AVAILABILITY STATEMENT

All data contained within the current manuscript stems from the utilization of novel code (provided via GitHub https://github.com/spechtk/SPECHTacle) in conjunction with publicly available, open-source datasets generated previously.

## AUTHOR CONTRIBUTIONS

KMS, conceptualization, data curation, pipeline development, code creation, formal analysis, writing – original draft, writing – review and editing. HCS, conceptualization, pipeline development, writing – original draft, writing – review and editing. MJR, conceptualization, supervision, funding acquisition, writing – review and editing.

## References

1. Tanay A, Regev A. Scaling single-cell genomics from phenomenology to mechanism. Nature. 2017;541:331–338. doi: 10.1038/nature21350

2. Okaty BW, Sturrock N, Escobedo Lozoya Y, Chang Y, Senft RA, Lyon KA, Alekseyenko OV, Dymecki SM. A single-cell transcriptomic and anatomic atlas of mouse dorsal raphe Pet1 neurons. eLife. 2020;9:e55523. doi: 10.7554/eLife.55523

3. Schiebinger G, Shu J, Tabaka M, Cleary B, Subramanian V, Solomon A, Gould J, Liu S, Lin S, Berube P, et al. Optimal-Transport Analysis of Single-Cell Gene Expression Identifies Developmental Trajectories in Reprogramming. Cell. 2019;176:928–943.e922. doi: 10.1016/j.cell.2019.01.006

4. Chehimi SN, Crist RC, Reiner BC. Unraveling Psychiatric Disorders through Neural Single-Cell Transcriptomics Approaches. Genes (Basel). 2023;14. doi: 10.3390/genes14030771

5. Garrido AN, Machhar R, Cruz-Correa OF, Ganatra D, Crome SQ, Wither J, Jurisica I, Gladman DD. Single-cell RNA sequencing of circulating immune cells supports inhibition of TNFAIP3 and NFKBIA translation as psoriatic arthritis biomarkers. Front Immunol. 2025;16:1483393. doi: 10.3389/fimmu.2025.1483393

6. Carangelo G, Magi A, Semeraro R. From multitude to singularity: An up-to-date overview of scRNA-seq data generation and analysis. Front Genet. 2022;13:994069. doi: 10.3389/fgene.2022.994069

7. Qin Y, Liu Z, Gao S, Martinez-Vasallo C, Long Y, Zhu X, Liu B, Gao Y, Xu X, Nohales MA, et al. 48-Hour and 24-Hour Time-lapse Single-nucleus Transcriptomics Reveal Cell-type specific Circadian Rhythms in Arabidopsis. Nat Commun. 2025;16:4171. doi: 10.1038/s41467-025-59424-8

8. Butto T, Mungikar K, Baumann P, Winter J, Lutz B, Gerber S. Nuclei on the Rise: When Nuclei-Based Methods Meet Next-Generation Sequencing. Cells. 2023;12. doi: 10.3390/cells12071051

9. Han L, Liu Z, Jing Z, Liu Y, Peng Y, Chang H, Lei J, Wang K, Xu Y, Liu W, et al. Single-cell spatial transcriptomic atlas of the whole mouse brain. Neuron. 2025;113:2141–2160 e2149. doi: 10.1016/j.neuron.2025.02.015

10. Wang C, Acosta D, McNutt M, Bian J, Ma A, Fu H, Ma Q. A single-cell and spatial RNA-seq database for Alzheimer’s disease (ssREAD). Nat Commun. 2024;15:4710. doi: 10.1038/s41467-024-49133-z

11. Wei Y, He Y, Liu Y, Zhu L, Feng T, Shen Z, Wei W, Liu L, Han L, Wang L. Gene expression profiling enables refined parcellation of cortical layers in the heterogeneous human cerebral cortex. Genome Med. 2026;18. doi: 10.1186/s13073-026-01704-z

12. He S, Li C, Lu M, Lin F, Hu S, Zhang J, Peng L, Li L. Comprehensive analysis of scRNA-seq and bulk RNA-seq reveals the non-cardiomyocytes heterogeneity and novel cell populations in dilated cardiomyopathy. J Transl Med. 2025;23:17. doi: 10.1186/s12967-024-05983-1

13. Kalucka J, de Rooij L, Goveia J, Rohlenova K, Dumas SJ, Meta E, Conchinha NV, Taverna F, Teuwen LA, Veys K, et al. Single-Cell Transcriptome Atlas of Murine Endothelial Cells. Cell. 2020;180:764–779 e720. doi: 10.1016/j.cell.2020.01.015

14. Li J, Wang J, Ibarra IL, Cheng X, Luecken MD, Lu J, Monavarfeshani A, Yan W, Zheng Y, Zuo Z, et al. Single-cell atlas of the transcriptome and chromatin accessibility in the human retina. Nat Genet. 2026;58:418–433. doi: 10.1038/s41588-025-02454-1

15. Theodoris CV, Xiao L, Chopra A, Chaffin MD, Al Sayed ZR, Hill MC, Mantineo H, Brydon EM, Zeng Z, Liu XS, et al. Transfer learning enables predictions in network biology. Nature. 2023;618:616–624. doi: 10.1038/s41586-023-06139-9

16. Chen H, Venkatesh MS, Gomicronmez Ortega J, Mahesh SV, Nandi TN, Madduri RK, Pelka K, Theodoris CV. Scaling and quantization of large-scale foundation model enables resource-efficient predictions in network biology. Nat Comput Sci. 2026;6:450–463. doi: 10.1038/s43588-026-00972-4

17. Gulati GS, D’Silva JP, Liu Y, Wang L, Newman AM. Profiling cell identity and tissue architecture with single-cell and spatial transcriptomics. Nat Rev Mol Cell Biol. 2025;26:11–31. doi: 10.1038/s41580-024-00768-2

18. Bard JBL. C.H. Waddington’s lasting achievements in development and evolution. Development. 2026;153. doi: 10.1242/dev.205836

19. Monge G. Memoire sur la theorie des deblais et des remblais. Mem Math Phys Acad Royale Sci. 1781:666–704.

20. Kantorovich L. On the transfer of masses (in russian). 1942.

21. Institute B. Tutorial (wot). https://broadinstitute.github.io/wot/tutorial/.

22. Cheng YC, Zhang Y, Tripathi S, Harshavardhan BV, Jolly MK, Schiebinger G, Levine H, McDonald TO, Michor F. Reconstruction of single-cell lineage trajectories and identification of diversity in fates during the epithelial-to-mesenchymal transition. Proc Natl Acad Sci U S A. 2024;121:e2406842121. doi: 10.1073/pnas.2406842121

23. Traxler L, Lucciola R, Herdy JR, Jones JR, Mertens J, Gage FH. Neural cell state shifts and fate loss in ageing and age-related diseases. Nat Rev Neurol. 2023;19:434–443. doi: 10.1038/s41582-023-00815-0

24. Schiebinger G, Shu J, Tabaka M, Cleary B, Subramanian V, Solomon A, Gould J, Liu S, Lin S, Berube P, et al. Optimal-Transport Analysis of Single-Cell Gene Expression Identifies Developmental Trajectories in Reprogramming. Cell. 2019;176:928–943 e922. doi: 10.1016/j.cell.2019.01.006

25. Vajapeyam S. Understanding Shannon’s Entropy metric for Information. 2014.

26. Ameri AJ, Lewis ZA. Shannon entropy as a metric for conditional gene expression in Neurospora crassa. G3 (Bethesda). 2021;11. doi: 10.1093/g3journal/jkab055

27. Teschendorff AE, Enver T. Single-cell entropy for accurate estimation of differentiation potency from a cell’s transcriptome. Nat Commun. 2017;8:15599. doi: 10.1038/ncomms15599

28. Grun D, Muraro MJ, Boisset JC, Wiebrands K, Lyubimova A, Dharmadhikari G, van den Born M, van Es J, Jansen E, Clevers H, et al. De Novo Prediction of Stem Cell Identity using Single-Cell Transcriptome Data. Cell Stem Cell. 2016;19:266–277. doi: 10.1016/j.stem.2016.05.010

29. Satija R, Farrell JA, Gennert D, Schier AF, Regev A. Spatial reconstruction of single-cell gene expression data. Nat Biotechnol. 2015;33:495–502. doi: 10.1038/nbt.3192

30. Choi JS, Ayupe AC, Beckedorff F, Catanuto P, McCartan R, Levay K, Park KK. Single-nucleus RNA sequencing of developing superior colliculus identifies neuronal diversity and candidate mediators of circuit assembly. Cell Rep. 2023;42:113037. doi: 10.1016/j.celrep.2023.113037

31. Wolf FA, Angerer P, Theis FJ. SCANPY: large-scale single-cell gene expression data analysis. Genome Biol. 2018;19:15. doi: 10.1186/s13059-017-1382-0

32. Shannon CE. A mathematical theory of communication. The Bell System Technical Journal. 1948;27:379–423. doi: 10.1002/j.1538-7305.1948.tb01338.x

33. Liao C, O’Farrell E, Qalieh Y, Savalia NK, Girgenti MJ, Kwan KY, Kwan AC. Single-nucleus transcriptomics reveals time-dependent and cell-type-specific effects of psilocybin on gene expression. bioRxiv. 2025. doi: 10.1101/2025.01.04.631335

34. Zheng R, Xu Z, Zeng Y, Wang E, Li M. SPIDE: A single cell potency inference method based on the local cell-specific network entropy. Methods. 2023;220:90–97. doi: 10.1016/j.ymeth.2023.11.006

